# A multimodal neuroprosthetic interface to record, modulate and classify electrophysiological correlates of cognitive function

**DOI:** 10.1101/2021.07.29.454271

**Authors:** Bettina Habelt, Christopher Wirth, Dzmitry Afanasenkau, Lyudmila Mihaylova, Christine Winter, Mahnaz Arvaneh, Ivan R. Minev, Nadine Bernhardt

## Abstract

Most mental disorders are characterised by impaired cognitive function and behaviour control. Their often chronic reoccurring nature and the lack of efficient therapies necessitate the development of new treatment strategies. Brain-computer interfaces, equipped with multiple sensing and stimulation abilities, offer a new toolbox, whose suitability for diagnosis and therapy of mental disorders has not yet been explored. Here, we developed a soft and multimodal neuroprosthesis to measure and modulate prefrontal neurophysiological features of neuropsychiatric symptoms. We implanted the device epidurally above the medial prefrontal cortex of rats and obtained auditory event-related brain potentials reflecting intact neural stimulus processing and alcohol-induced neural impairments. Moreover, implant-driven electrical and pharmacological stimulation enabled successful modulation of neural activity. Finally, we developed machine learning algorithms which can deal with sparsity in the data and distinguish effects with high accuracy. Our work underlines the potential of multimodal bioelectronic systems to enable a personalised and optimised therapy.

## Introduction

The prefrontal cortex (PFC) represents a fundamental structure for behavioural top-down control and modulates attention, working memory, reward evaluation and the ability to self-control actions, emotions and stress. Disturbances of prefrontal neural network activity underlie many cognitive and behavioural impairments observed in neuropsychiatric diseases such as addictive disorders, schizophrenia or autism. The complex aetiology of these diseases makes optimal treatment challenging (*1*). Current therapeutical interventions often present with side effects and lack long-term efficacy. In particular, systemic pharmacotherapy shows poor topical specificity and adaptability to changes in patient-specific needs regarding treatment duration and intensity (*2*). The chronic reoccurrence and high relapse rates following treatment (*3*) thus warrant the development of new treatment approaches.

Brain stimulation techniques, such as Deep Brain Stimulation (DBS) and transcranial Direct Current Stimulation (tDCS), have gained increasing attention as alternative treatment options. DBS can be precisely delivered to specific brain areas through deeply implanted electrodes. Predominantly applied in Parkinson’s disease to improve motor function (*4*), DBS also showed beneficial effects in neuropsychiatric disorders such as depression (*5*), drug addiction (*6, 7*), as well as obsessive compulsive- and anxiety disorders (*8*). However, due to its invasiveness and continuous stimulation mode, DBS holds the risk of side effects such as impaired speech, gait and cognition (*9*) and has therefore been restricted to a small number of severe and otherwise treatment resistant cases. In contrast, tDCS offers scalp-applied and thus non-invasive brain stimulation with no or just minimal side effects (*10*). Prefrontal tDCS has been shown to reduce symptoms of depression (*11*) and schizophrenia (*12*) as well as craving and drug consumption in substance use disorders (*13*). However, reports of varying treatment efficacy (*14*) in response to tDCS might be due to identical stimulation parameters used for all subjects in a rigid “one-size-fits-all” fashion not taking into account individual differences in brain anatomy, underlying pathology and temporal changes in brain states (*15*). Furthermore, up to ∼75 % of epicranially applied currents are attenuated by scalp and skull (*16*) hampering target region and dose specification.

Epicortical neuroprosthetics, equipped with multiple sensing and stimulation abilities, might offer a new off-the-beaten-track toolbox for diagnosis and therapy and may overcome some of the limitations of current brain stimulation techniques. Implanted epi- or subdurally and made of soft and biocompatible materials (*17*–*19*), epicortical devices can adapt to the curvilinear surface of the brain resulting in reduced tissue inflammation and improved long-term stability compared to brain penetrating electrodes (*20, 21*). Furthermore, direct cortical stimulation via small surface electrodes provides effective and precise stimulation close to the target structure (*22*). Furthermore, epicortical neuroprosthesis enables combined neuromonitoring and stimulation in one device allowing immediate detection of stimulation effects on neural activity. Current clinical applications of epicortical electrodes for electrocorticography (ECoG) and direct cortical stimulation focus on real-time functional brain mapping to assess language, motor and sensory function during surgical intervention for medically intractable epilepsy and brain tumours (*22, 23*). Besides intraoperative epileptic seizure localisation, an ECoG-type array combined with direct cortical stimulation has been successfully implemented to reduce an incipient seizure by detecting abnormal neural activity that subsequently triggers stimulation. This so-called Responsive Neurostimulation System^®^ (NeuroPace^®^, Mountain View, CA, U.S.A.) is the first demonstration of a genuinely bidirectional, closed-loop brain-computer interface approved for clinical application (*22, 24*).

Closed-loop neuromodulation may also open opportunities in the treatment of neuropsychiatric disorders by utilising artificial intelligence and advanced machine learning algorithms to identify brain states and optimise stimulation parameters on the basis of pre-defined neural features (*25*). Event-related potentials (ERPs) present eligible biomarkers for this purpose. These short electrical deflections induced in the brain immediately following an external or internal event have proven valuable for investigating sensory information processing and higher-order cognition in healthy individuals as well as psychopathological conditions (*26*). Within scalp-recorded electroencephalograms (EEG), ERPs appear as time-locked local negative or positive maxima of a few microvolts (µV) lasting tens to hundreds of milliseconds (ms). They are commonly labelled according to their polarity (negative = N, positive = P) and latency (in ms post-stimulus or order of appearance within the recorded waveform) (*27*). The best-known paradigm to elicit ERPs is the “oddball” paradigm in which subjects are confronted with a series of frequent (e.g. auditory or visual) stimuli (”standards“) randomly interspersed with rare stimuli (_”_deviants“). Common ERP components observed in such tasks include P1, N1, P2, N2 and P3. The components P1, N1 and P2 reflect pre- and early attentive automatic stimulus processing and sensory gating that constitutes an inhibitory filter mechanism to enable focusing on salient stimuli while disregarding irrelevant or repetitive information (*28*). The N2 is elicited by rare events and reflects a change-detection response sensitive to novelty and stimulus probability (*29, 30*). Likewise, amplitudes of the P3 vary with stimulus incidence and significance but also depend on a subject’s motivation, attentional resources and cognitive capabilities (*29, 31*). Modified ERPs indicate impaired PFC functioning and cognitive deficits associated with neuropsychiatric diseases (*32*). For example, delayed and/or reduced ERP amplitudes have been observed in alcohol-addicted patients and animal models (*33*–*36*). Primarily a disturbed P3 component has been related to poor behaviour control and increased relapse probability and therefore judged as a suitable predictor for the relapse risk after drug withdrawal (*37, 38*).

ERPs measured by scalp-EEG have high temporal precision but lack spatial resolution, are sensitive to noise, and, like in tDCS, electrical signals are partly silenced through the skull (*39*). ECoG electrodes are in closer proximity to the source of relevant brain activity and have demonstrated superior signal sensitivity, broader bandwidth, higher topographical resolution and a lower vulnerability to artefacts than EEG resulting in accurate ERP acquisition (*21, 40*).

So far, ECoG electrodes have been placed over lateral, sensorimotor areas based on clinical requirements for epileptic seizure localisation (*40, 41*) or to enable paralysed patients to control external devices using movement-related neural activity patterns (*42, 43*). However, the potential of an epicortical implant to target cognitive ERPs from central prefrontal brain regions remains unexplored.

Based on the advantages and opportunities of an implanted bidirectional brain-computer interface, we set out to build a tailor-made soft and multimodal epicortical device to measure and modulate neurophysiological features relevant to the diagnosis and therapy of neuropsychiatric disorders. We implanted our device epidurally above the medial (m) PFC of rats and tested its feasibility to obtain auditory ERPs. We detected activity alterations induced by acute alcohol intake and implant-driven neuromodulation with direct application of electrical pulses and pharmacoactive naltrexone (NTX). We furthermore deployed machine learning algorithms to distinguish treatment-specific brain responses from single ERP trials with potential use as feedback for closed-loop adjustment of neurostimulation in a therapeutic neuroprosthesis. Finally, we performed an immunohistochemical analysis of implant and intervention tolerability.

## Results

### Development and implantation of an epicortical neuroprosthesis to interface the PFC

We initially developed a custom implantable device covering the surface of the frontal lobe of the rat cortex. Implants consisted of electrodes arranged in a 3 × 3 matrix and labelled according to their position above the mPFC as frontocentral (FC), frontal left (FL), frontal right (FR), medial central (MC), medial left (ML), medial right (MR), posterior central (PC), posterior left (PL) and posterior right (PR). Eight of the electrodes (0.2 × 0.2 mm^2^) were used for neural recording only. The larger FC electrode (1 × 1 mm^2^) was used for both, recording and electrical stimulation of the mPFC spanning both hemispheres at 3.2 mm anterior to bregma. A microfluidic channel was integrated to enable local delivery of liquids at 2.2 mm anterior to bregma. We used a recently established prototyping technology allowing rapid fabrication of soft and customised bioelectronic implants. Thereby, arrays were 3D-printed layer-by-layer using soft silicones and a conductive platinum ink (Fig. 1 A-C) (*17, 19*).

**Fig. 1.**
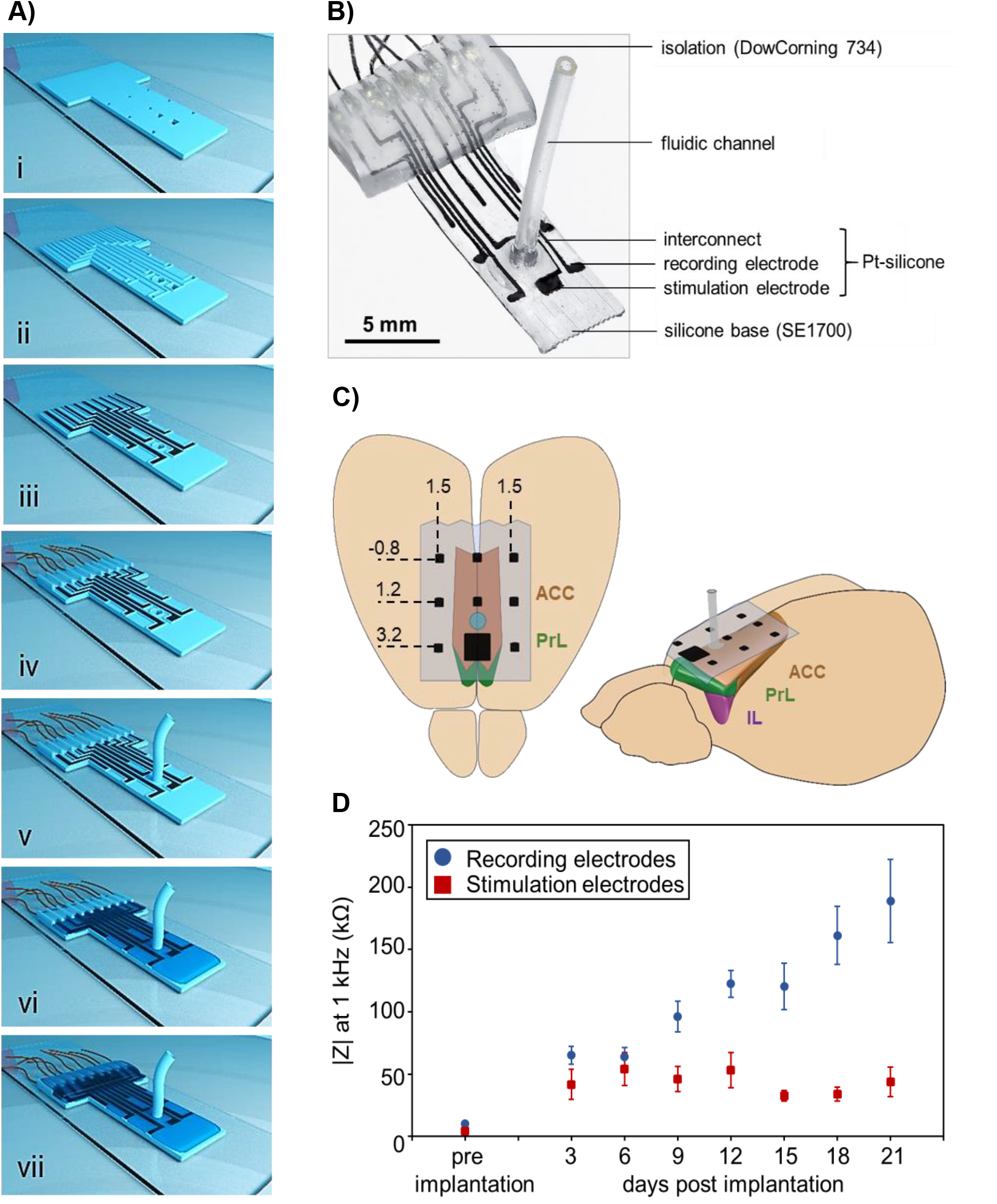
Multimodal epicortical array for recording and modulation of neural activity. **(A)** i) Silicone base layer (DOWSIL™ SE1700) including holes defining the active sites of electrodes and the microfluidic channel. ii) A second silicone layer defines the borders for electrode interconnects and the microfluidic channel. iii) The conductive portions of the array are ink-jet printed from platinum-silicone composite. iv) Connection of microwires. v) Connection of silicone fluidic channel. vi) Isolation of the array using SYLGARD™184. vii) Isolation of the cable contacts using DOW CORNING^®^ 734. **(B)** Photograph of an implant **(C)** Implantation of the array above the rat mPFC encompassing anterior cingulate (ACC), prelimbic (PrL) and infralimbic (IL) cortices. Stereotactic coordinates (in millimeters) are relative to bregma. **(D)** Impedances *in vitro* (at 1 kHz) of recording (blue, n = 80 electrodes of 10 implants) and stimulation electrodes (red, n = 10 electrodes of 10 implants) and *in vivo* (varying implant numbers, see Table S 1). Data are presented as mean ± SEM.

Electrode impedances at 1 kHz measured *in vitro* were 10.14 ± 1.96 kΩ (mean ± standard error of the mean (SEM)) for recording electrodes (n = 80 from 10 implants) and 4.36 ± 1.41 kΩ (n = 10 from 10 implants) for stimulating electrodes (Fig. 1 D, Fig. S 1, Table S 1).

The bioelectronic devices were implanted epidurally above the mPFC (Fig. 1 C) of 10 rats in a delicate surgical procedure involving trepanation directly above the superior sagittal sinus and adjacent blood vessels. Attention was paid to limit drilling to a few seconds at a time at a low drill rotational speed under constant flushing with cold phosphate-buffered saline (PBS) to prevent thermal tissue damage though occasional microbleedings could not be avoided. To allow influx of NTX solution, the dura was incised bilaterally about 0.5 mm next to the position of the microfluidic channel.

The stability of implants *in vivo* was evaluated for up to 3 weeks, during which impedances of individual electrodes were measured every third day. A general trend of impedance increase was observed over time in recording electrodes, while impedances of stimulating electrodes remained stable (Fig. 1 D, Table S 1). Functional electrodes suitable for further ERP analysis were defined as having an impedance below 600 kΩ. Throughout the study period, more than 80 % of electrodes remained functional (Table S 1). One of the stimulation electrodes lost functionality at the last session time point.

### Implementation of ECoG measurements of prefrontal cortical activity

We set out to establish if our technology is capable to reliably obtain characteristic ERPs in awake rats (n = 10). We used an auditory oddball paradigm similar to those applied in human ERP studies to elicit typical sound-specific ERP responses. To reduce movement artefacts, rats were placed in a sling within an electrically shielded and soundproofed audiometry booth (Fig. 2). During a 5 × 5 min session, rats passively listened to a series of frequent standard sounds (50 ms, 1 kHz, 70 dB sound pressure level (SPL), 1200 times = 87 % of trials) presented once every second and randomly interspersed with rare deviant sounds (50 ms, 2 kHz, 80 dB SPL, 180 times = 13 % of trials).

**Fig. 2.**
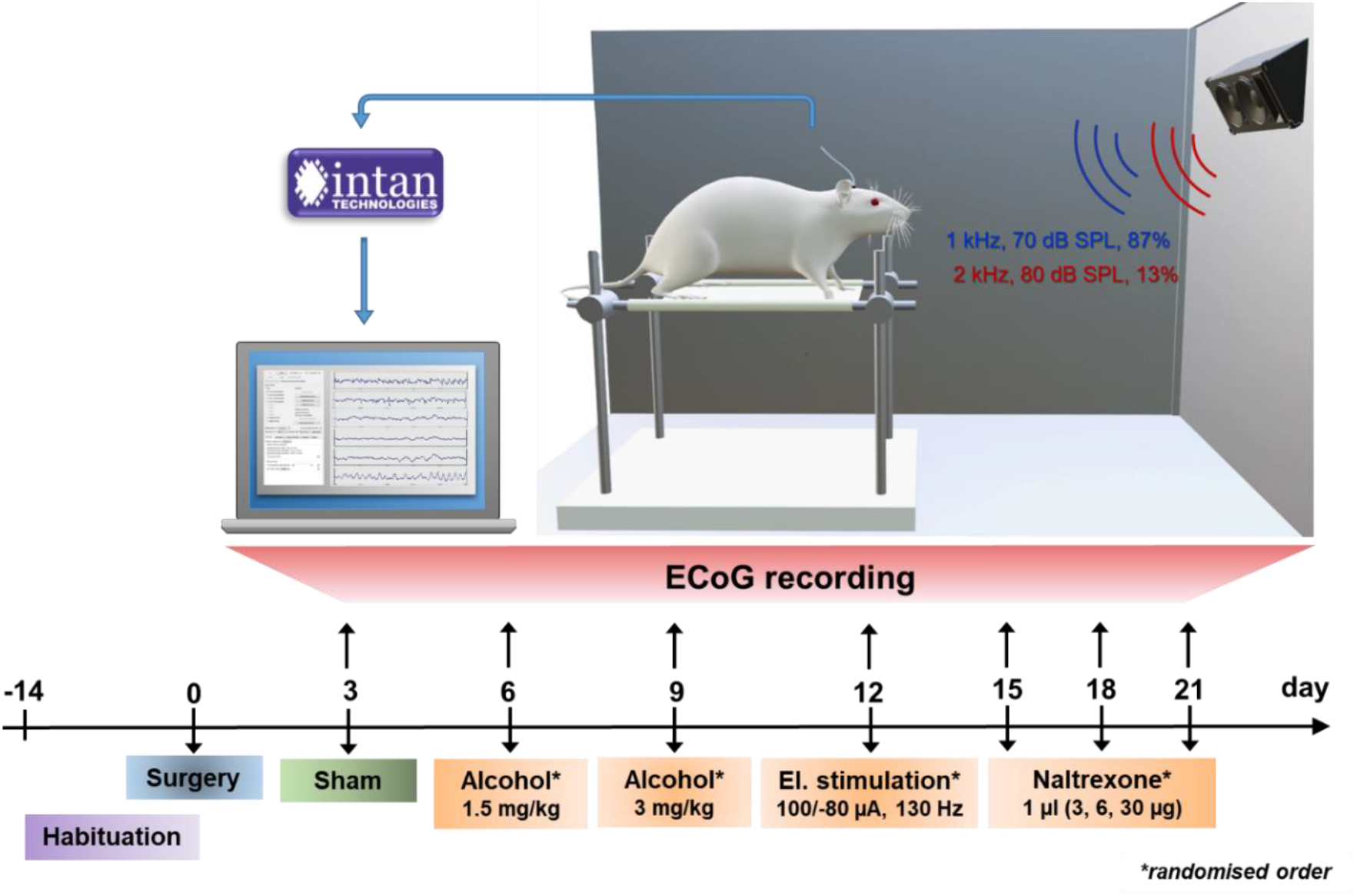
Set-up and timeline for ECoG recording sessions. Experiments were planned as a within-subjects-design with all animals undergoing an initial habituation period, surgical intervention and neural recordings without previous treatment (sham) and following alcohol injections, electrical brain stimulation and cortical administrations of NTX in a randomised order. Electrical potentials elicited during the auditory oddball task were amplified and digitised at a sampling rate of 3 kHz via the Intan RHD2000 recording system and visualised and saved on a computer using the Intan recording software.

The acquired neural data were initially pre-processed involving filtering, data segmentation and artefact rejection followed by averaging standard and deviant trials for each animal. Then, ERP amplitudes were computed as averages within a time window of 10 ms around the peak latency determined within the following time intervals post-stimulus: P1: 30 – 75 ms, N1: 80 – 105 ms, P2: 110 – 125 ms, N2: 130 – 180 ms, P3: 200 – 500 ms. As the positive P2 component was observed to be in the negative value range here, the N1-P2 peak-to-peak difference was exceptionally used for calculating the P2 amplitude.

Differences in neural activity underlying perception of the standard and deviant sounds should result in different voltage waveforms recorded through the implant. To determine if the neural responses to the two sounds can be distinguished in the ECoG recordings, we performed statistical analysis using a channel-wise one-sample *t*-test of the difference curve (deviant minus standard) against zero value. The resulting *p*-values from these contrasts were corrected post-hoc for multiple comparisons and additionally reported as FDR (false discovery rate)-adjusted *p*-values. Effect sizes were calculated using Cohen’s *d* with |*d*| ≥ 0.2 indicating a small, |*d*| ≥ 0.5 a medium and |*d*| > 0.8 a large effect.

Grand average ERPs over all animals revealed significantly different neural activities elicited by the two sounds with large effect sizes for P1, N1 and P3 components at all electrode sites, also bearing FDR correction (Fig. 3, Table S 2). The components P2 and N2 were not clearly recognisable at each electrode though significant differences within these time intervals have been detected at some channels (e.g. electrode FL, P2: *t*(9) = 2.318, *p* = 0.046 uncorrected; N2: *t*(9) = -2.514, *p* = 0.033 uncorrected) but did not withstand FDR correction.

**Fig. 3.**
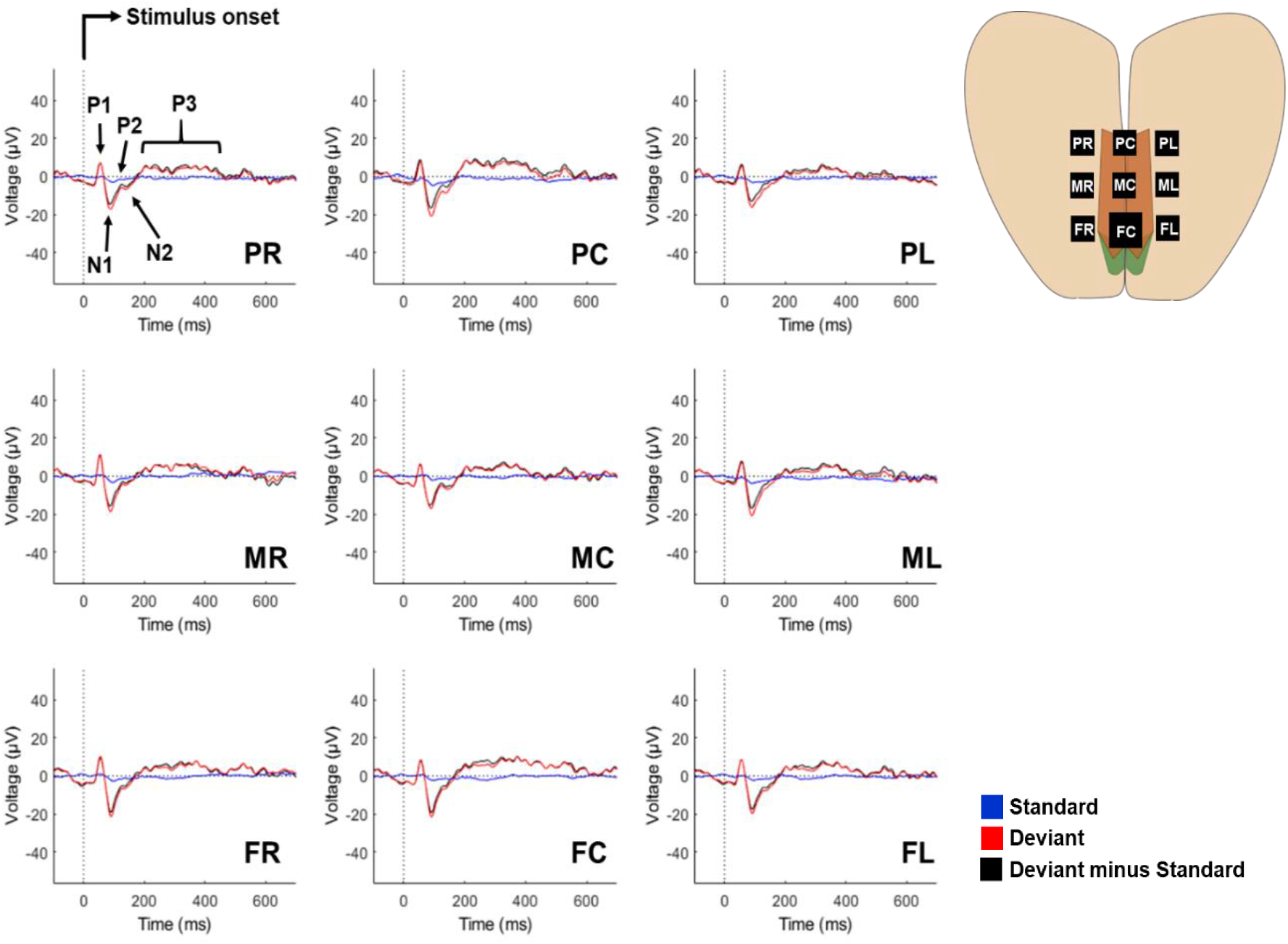
Grand average ERPs. elicited by the standard sound (blue), deviant sound (red) and their difference curves (black) at all electrode sites. Traces are averaged from 10 animals. The components P1-N1-P2 are characteristic for pre- and early attentive auditory signal processing while N2 and P3 are indicative for conscious stimulus evaluation.

### Electrical and pharmacological modulation of neural activity

Following the successful acquisition of ERPs in untreated animals (sham), we tested if we can detect changes in neural activity induced by acute systemic alcohol administration, as well as electrical stimulation and NTX, both applied directly to the cortex through the implant. Following each of the interventions, animals were subjected to the same auditory oddball task. Interventions could not be performed on all animals due to issues with connectors, cement adhesion or blockage of the microfluidic channel, which necessitated these animals to be dropped from the study.

In each treatment condition, neural responses to the standard sound appeared flat, as was the case for untreated animals (presumably due to habituation effects caused by the high rate of repetitions). Therefore and in order to focus on treatment-induced changes of neural activity, subsequent analysis was performed on ERP difference curves (deviant-minus-standard sounds).

Animals received a low (1.5 g/kg, n = 6) or a high (3.0 g/kg, n = 9) dose of ethanol (EtOH, 20 % v/v, injected intraperitoneally, i.p) about 20 min before recording. Behaviourally, the low dose induced a slightly tottering gait, while the high dose resulted in ataxia and immobility. In line, paired *t*-tests revealed a delay of the N2 component at four channels in the low ethanol condition, but no significant impact on ERP amplitudes (Fig. 4, Fig. S 2, Table S 3, S 4). In contrast, high dosed ethanol significantly impaired neural functioning as reflected by a diminished N1 component (Fig. 4, Fig. S 2, Table S 6). Analysis of ERP latencies for high ethanol was performed for the P1 component (Table S 6) as other ERP components were suppressed entirely.

**Fig. 4.**
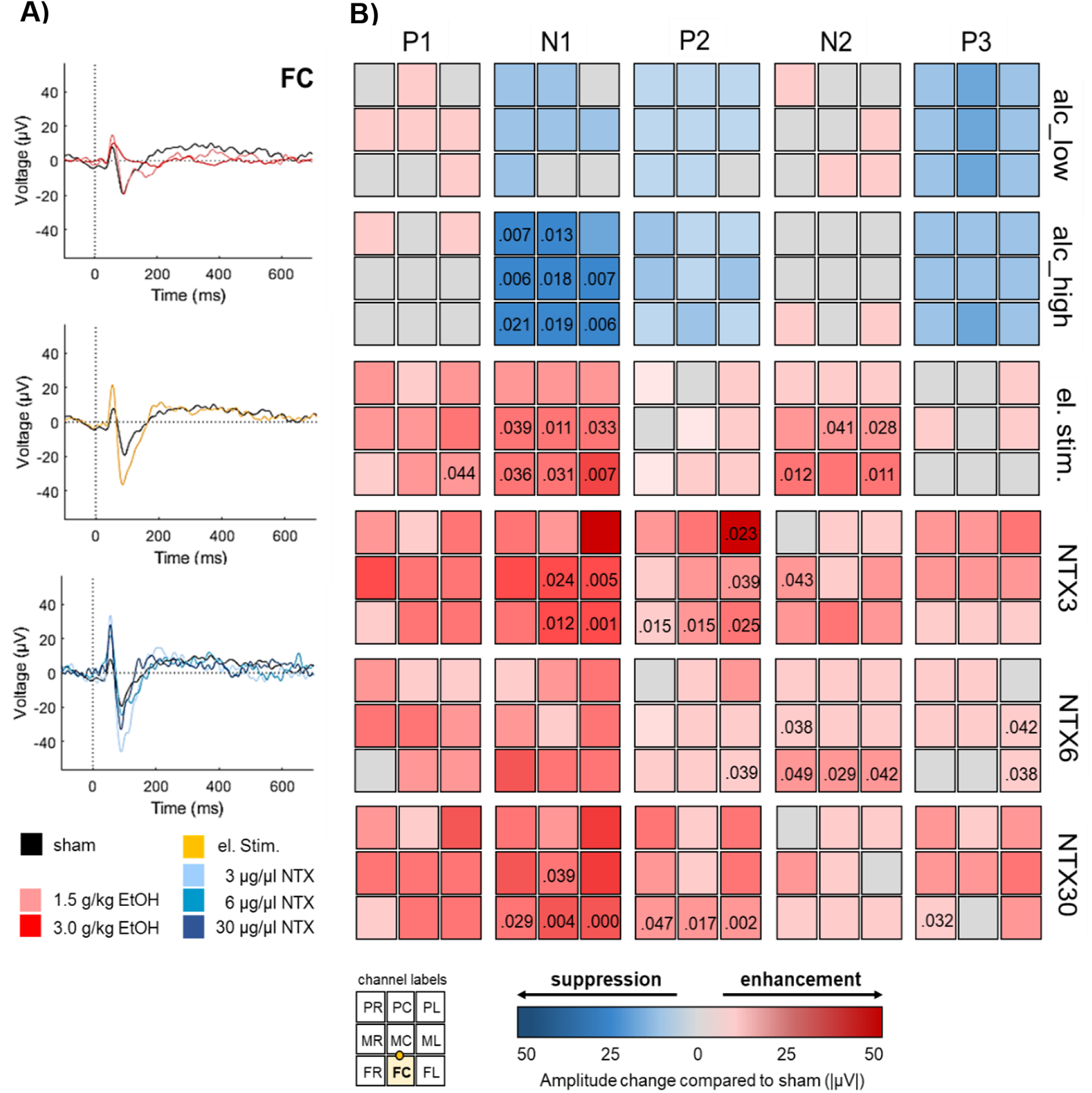
Impact of alcohol and implant-driven electrical and pharmacological brain stimulation on prefrontal neural activity. **(A)** Representative grand average deviant-minus-standard ERP difference curves at channel FC following administration of alcohol at 1.5 g/kg (n = 6, rose) and 3 g/kg (n = 9, red), electrical brain stimulation (n = 8, yellow), cortical delivery of naltrexone (NTX) at 3 µg/µl (n = 5, light blue), 6 µg/µl (n = 4, mid-blue) and 30 µg/µl (n = 5, dark blue) and untreated animals (sham, n = 10, black). **(B)** ERP amplitude differences between interventions and untreated animals at all channels. Implant channels are displayed in 3 × 3 matrices for each treatment (rows) and ERP component (columns) with suppressing or enhancing treatment effects on ERP amplitudes illustrated in blueish or reddish colours, respectively. The bottom left inset illustrates channel locations with the highlighted FC stimulation electrode and the microfluidic channel between electrodes FC and MC. Numbers indicate significant p-values (uncorrected).

Next, we investigated the effects of direct cortical stimulation on neural activity by applying biphasic, charge-imbalanced pulses (20 min, 100 µA anodal/ 80 µA cathodal, 130 Hz) as such waveforms have been shown to provide a good compromise between effective neural activation and adverse effects such as tissue damage and dissolution of platinum electrodes (*44*). Animals (n = 8) did not display behavioural changes upon stimulation, however increased P1, N1 and/or N2 amplitudes at six channels indicated an enhancing effect of electrical stimulation on brain activity (Fig. 4, Fig. S 3, Table S 8). For the N1 component, this effect was more pronounced in closer proximity to the stimulation site as revealed by paired *t*-tests using mean N1 responses combined for frontal (*t*(6) = -4.703, *p* = 0.003, |*d*| = 0.936) and mid row electrodes (*t*(6) = -3.139, *p* = 0.020, |*d*| = 0.516) compared to posterior electrode sites (Fig. 4 B). ERP latencies were unchanged (Table S 7).

Finally, we investigated the effects of epicortical administration of NTX, an opioid receptor antagonist well established in recuperation but with just moderate effects in conventional oral application (45). We tested three different doses (3 µg (NTX3, n = 5), 6 µg (NTX6, n = 4), 30 µg (NTX30, n = 5)), dissolved in artificial cerebrospinal fluid and administered in a volume of 1 µl via the integrated microfluidic channel. Differences in animal behavior upon NTX administration at either of the tested doses were not observed. NTX at 3 µg/µl and 30 µg/µl decreased latencies of P1 and N1 components (Table S 9, S 13) and all concentrations displayed enhancing effects on amplitudes of N1, P2, N2 or P3 components (Fig. 4, Fig. S 4, Table S 10, S 12, S 14). However, results remained significant following FDR-correction only at channels near the microchannel outlet for N1 amplitudes following 3 µg/µl (channels FC, MC, FL, ML) and 30 µg/µl (channels FC, FL) and for the P2 at channel FL following 30 µg/µl.

### Single-trial ERP classification

Computing ERP grand averages in standard offline ERP analysis procedures is unsuitable for a neuroprosthetic device designed to operate in real-time. Thus, we applied machine learning algorithms to perform single-trial classification of ERP responses. We aimed to see if the aggregate of the classifier outputs provided an accurate differentiation of which treatment had been applied to each session for each individual. Classifications were performed comparing one-vs-one combinations of all treatments as depicted in Table 1.

**Table 1.** Percentage of sessions in which the treatment was accurately predicted for each one-vs-one treatment type comparisons.

| Treatment A | Treatment B | n | Session Classification Accuracy | *p*-value |
| --- | --- | --- | --- | --- |
| sham | alcohol high | 9 | 94.4 % | < 0.001 |
| sham | alcohol low | 6 | 66.7 % | 0.248 |
| alcohol low | alcohol high | 6 | 83.3 % | 0.014 |
| sham | el. stimulation | 8 | 75.0 % | 0.046 |
| alcohol high | el. stimulation | 7 | 92.9 % | 0.001 |
| alcohol low | el. stimulation | 6 | 100.0 % | < 0.001 |
| sham | NTX3 | 4 | 87.5 % | 0.029 |
| sham | NTX6 | 4 | 75.0 % | 0.103 |
| sham | NTX30 | 5 | 90.0 % | 0.001 |
| alcohol high | NTX3 | 3 | 100.0 % | 0.014 |
| alcohol high | NTX6 | 3 | 83.3 % | 0.083 |
| alcohol high | NTX30 | 4 | 75.0 % | 0.103 |
| alcohol low | NTX3 | 3 | 66.7 % | 0.410 |
| alcohol low | NTX6 | 3 | 66.7 % | 0.410 |
| alcohol low | NTX30 | 4 | 75.0 % | 0.103 |

We used the FC electrode as this was the channel in which successful recordings were consistently available, only missing NTX3 for one animal. Furthermore, specific ERP components, such as N1 and P3, are known to peak at frontocentral electrode sites (*29, 46*). Data were low-pass filtered and resampled to 64 Hz to reduce dimensionality, thus decreasing the possibility of overfitting (*47*). As grand average data showed notable differences between sham recordings and EtOH treatments, especially of the N1 and P3, time-domain data in the time windows of these components were chosen for feature extraction based on voltage differences between sham recordings and treatments. Next, ERP difference trials were generated as shown in Fig. 5 A to represent the contrast between responses to standard and deviant sounds in a given condition. To train a model predicting the treatment applied in a particular session for a given animal, all difference trials from all other animals were combined to form the training data. No trials from the test animal were used as training trials, as these could bias the results and reduce the generalisability of the training model. Difference trials from the test animal under a given treatment were used as the test data. Next, we performed a stepwise linear discriminant analysis (SWLDA), chosen as a combined feature selection and classification strategy. SWLDA has previously been proven effective in these areas when processing P3 data (*48*), including performing single-trial classification (*49*). Putative feature sets were generated to find the feature set that provided the best separability of the two classes of training data. Starting with an empty model, individual features were systematically added to or removed from the model at each step until no further improvements could be made to the model. At this stage, the features in the model were used as the selected set, as depicted in Fig. 5 B. Then, linear discriminant analysis involving the stepwise selected features was carried out to classify each test trial from a given session under either treatment A or B (Fig. 5 C). Finally, a simple majority vote predicted the overall treatment class of the session, as shown in Fig. 5 D.

**Fig. 5.**
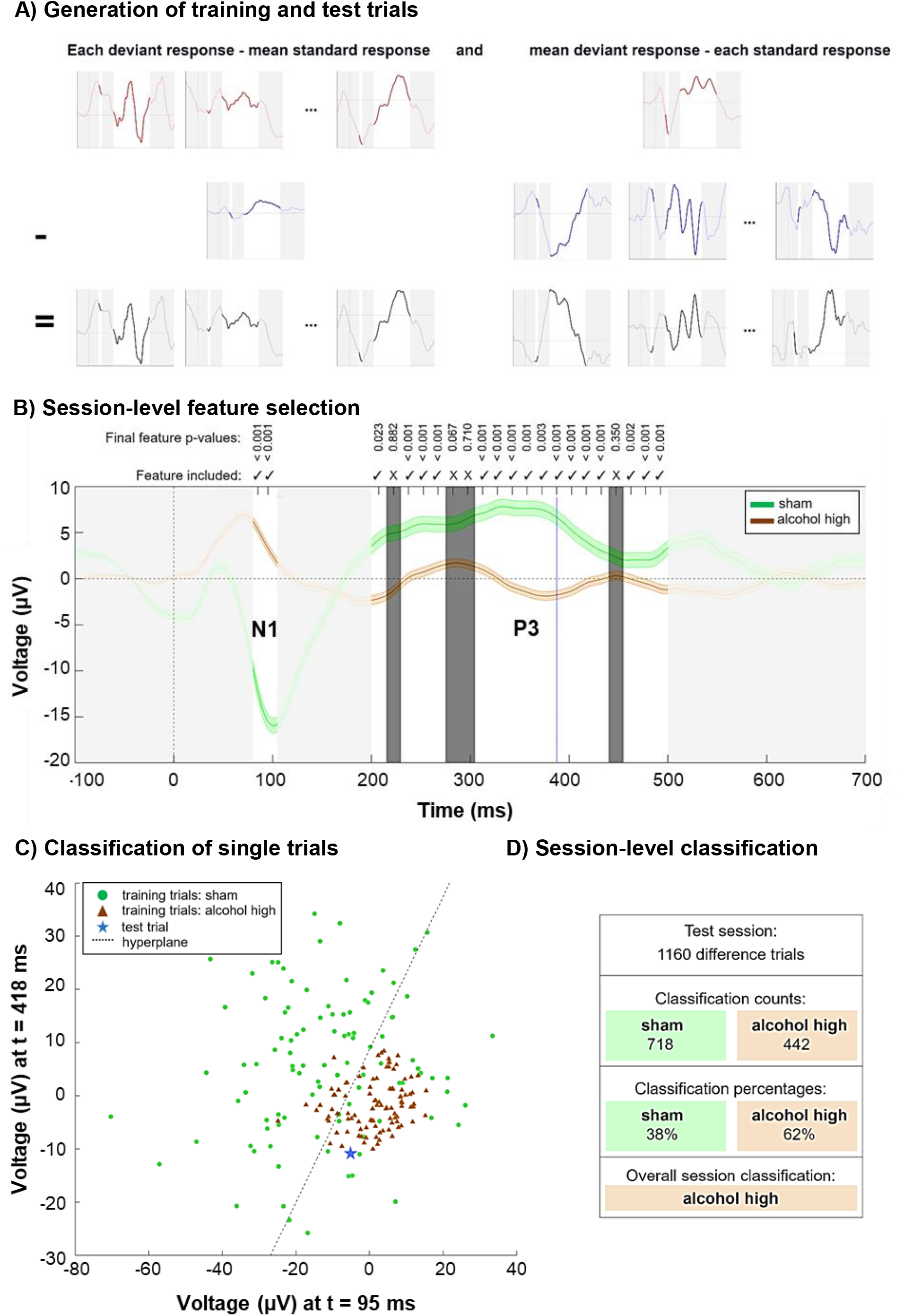
Approach to distinguish the treatment applied for each session. **(A)** Generation of ERP difference trials for a single animal and treatment. Upper row (red): deviant ERP responses, including the mean (right). Mid row: standard ERP responses, including the mean (left). Bottom row: resulting difference trials. Time ranges that were not used for classification are obscured in light grey. **(B)** Example result of the stepwise feature selection phase representing a model for testing whether the animal’s session could be correctly classified as ‘alcohol high’ against ‘sham’ treatment. Solid green line: mean of all difference trials under ‘sham’ treatment. Solid brown line: mean of all difference trials under ‘alcohol high’ treatment. Shaded areas above and below these lines represent 1 standard error. Each individual feature represents the voltage at a given time point of each trial. For example, the feature highlighted by the blue dotted line represents voltages of each trial at 388 ms post-stimulus. Time ranges, not eligible for classification, are obscured in light grey. Non-selected features are obscured in dark grey and marked with crosses above. Features selected for the classification model remain visible with check marks above. The *p*-values reported in the last iteration of stepwise feature selection are reported alongside the status of each feature as either included or excluded. **(C)** Simplified example of the linear discriminant classifier employed to classify each individual trial as a result from a specific treatment type. For the purpose of visualisation, the two features with smallest *p*-values in the feature selection example shown in part B are extracted and the 100 samples closest to the arithmetic mean of each class are displayed. Green circles: samples from ‘sham’ treatment. Brown triangle: samples from ‘alcohol high’ treatment. The hyperplane generated by the classification model best separating the two classes, is indicated as a dashed black line. An example test sample is shown as a blue star. As this sample falls on the lower right side of the hyperplane, it is correctly classified as an ‘alcohol high’ sample. **(D)** Overall classification made by the model. After each test trial (as generated in part A and tested as in part C with the features selected in part B), the majority of trials in this example was classified as ‘alcohol high’. Therefore, the overall prediction for the session is ‘alcohol high’.

The accuracy of the predictions for each one-vs-one comparison of treatment types is presented in Table 1. When comparing sham condition against treatment of 3 g/kg of EtOH, the correct treatment was predicted for 94.4 % of sessions, with only one misclassification. Furthermore, brain states induced by electrical stimulation were correctly distinguished from all other conditions with a precision up to 100 % (low EtOH dose). Pharmacological treatments with NTX were correctly classified in at least two-thirds of sessions for every comparison. Contingency tables reporting the predictions in each treatment comparison are shown in Supplementary Table S 15 – S 29.

### Biocompatibility of implant materials and applied treatments

Finally, we investigated the biointegration of the device itself and the effects of the combined interventions on brain tissue after four weeks of implantation. This involved immunohistochemical evaluation of neuroinflammation by applying antibodies against glial fibrillaric acidic protein (GFAP) and ionized calcium binding adaptor molecule 1 (Iba1) and stainings of laminins and platelet endothelial cell adhesion molecule (also known as cluster of differentiation 31 (CD31)), revealing cerebral vascular integrity. We furthermore investigated neuronal cell survival by using antibodies against the neuronal marker Hexaribonucleotide Binding Protein-3 (NeuN) and cysteine-aspartic protease (caspase3), a mediator of programmed cell death.

Immunostainings were performed on three experimental groups: 1.) sham-operated animals without implants (n = 3), 2.) animals that received a non-functional dummy implant (n = 3) and 3.) rats with an active implant that underwent EtOH injections, electrical stimulation and NTX delivery (n = 7). Each fluorescence staining was performed on six slices from each brain. For subsequent image analysis, we defined a region of interest (ROI) covering the entire implant width and the cortex up to a depth of 2 mm (Fig. 6 A). Zoomed-in microscopic images of Fig. 6 B depict the immunoreactivity within the left motor cortex at 3.2 mm anterior to bregma next to the stimulation electrode. The percentage of stained area, counts of stained objects per mm^2^ and mean fluorescence intensity were averaged over brain slices for each animal for statistical analysis (Fig. 6 C, Table S 30).

**Fig. 6.**
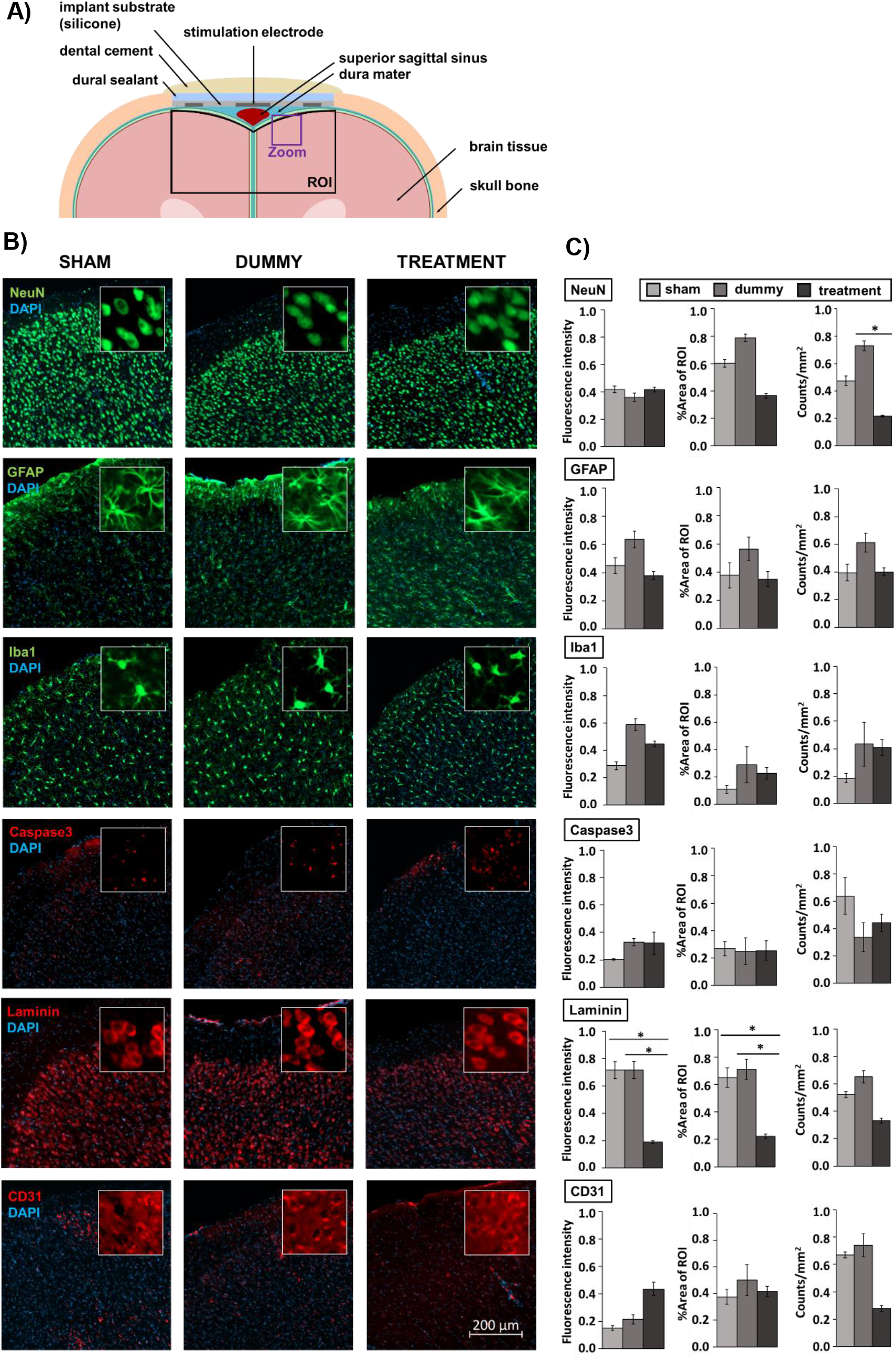
Immunohistochemical evaluation of implant and treatment tissue response. **(A)** Cross section of rat brain and the biomedical implant at 3.2 mm anterior to bregma including the region of interest (ROI) defined for biocompatibility analysis following device explantation and magnified view presented in (B). **(B)** Representative microscopic images of the left motor cortex at 3.2 mm anterior to bregma next to the stimulation electrode. Insets are 50 × 50 µm squares. **(C)** Selected biotolerance indicators for each, six brain slices per staining of sham-operated animals (n = 3), rats that received an implant dummy (n = 3) and treatments (n = 7) are normalised and presented as mean ± SEM. Asterisks indicate significant differences (*p* < 0.05).

In line with other studies using ECoG implants (*20, 50*), we observed fibrous tissue surrounding the electrode array leading to a slight depression of brain tissue underneath. We observed a lower number of NeuN positive cells in the treatment group (*F*(2,10) = 4.474, *p* = 0.045) indicative for an effect of treatment rather than caused by the device as animals that received an implant dummy did not display a significant difference in %area stained and numbers of NeuN positive cells compared to sham-operated rats (Fig. 6 B, C, Table S 30). No differences between groups were observed for GFAP, Iba1and caspase3, indicating that neither the dummy implant nor the treatments induced significantly enhanced inflammation or acute cell loss (Fig. 6 B, C, Table S 30). However, we observed treatment-related cellular alterations within the cerebral vascular system indicated by lower laminin expression in animals of the treatment group compared to sham-operated rats (fluorescence intensity: *F*(2,10) = 8.793, *p* = 0.021, %area of ROI: *F*(2,10) = 7.853, *p* = 0.042) and rats with implant dummies (fluorescence intensity: *F*(2,10) = 8.793, *p* = 0.021, %area of ROI: *F*(2,10) = 7.853, *p* = 0.020) (Fig. 6 B, C, Table S 30)).

## Discussion

We established a multimodal neural interface in combination with machine learning to acquire, modulate and classify ERP components in awake rats. While available ECoG-based systems typically focus on sensorimotor brain areas, this study proposes the first-time application of an epicortical implant to target cognitive ERPs from higher-order prefrontal networks. The applied tools have potential applications in diagnosing and treating neuropsychiatric diseases and aim to pave the way for intelligent closed-loop neuroprostheses.

Custom electrode arrays were manufactured using a 3D printing technology with robot-controlled deposition of soft and conductive materials that allowed tailoring the implant layout to match the surface area above the mPFC. The flexible Pt-silicone electrodes of our arrays provided low impedances *in vitro*. Previous studies demonstrate that electrode impedances increase during the first weeks after implantation but decline over more extended periods (*51*). Similarly, we observed increasing impedances at 1 kHz of recording electrodes during the 3-week study period though we did not perform measurements over longer implantation durations to clarify if a decrease in impedances would also occur in our devices. Impedances of stimulation electrodes were lower compared to recording electrodes due to their larger surface area and remained stable over the entire study period. A potential explanation for this may be that electrical stimulation supposedly has a “rejuvenating” effect on the electrode-tissue interface by decreasing the tissue interface resistance resulting in improved signal quality and reduced electrode impedances (*52*). The stability enabled reliable and high-quality field potential recordings.

Neuropsychiatric disorders are associated with disturbances in prefrontal brain activity that also translate into altered ERPs. Their monitoring is therefore increasingly supported to become part of the clinical routine (*53*). We recorded ERPs appearing in the brain as early stages of auditory perception immediately after the onset of sounds. In healthy subjects, they differ depending on sound characteristics (e.g. pitch, loudness). Our bioelectronic implant was capable to reliably measure these ERP differences between rare deviant and frequent standard sounds. Likewise, we could successfully detect dose-dependent ERP changes following acute administration of alcohol. In line with previous studies in humans and rodents, the application of EtOH substantially affected the N1 component indicating impaired sensory gating and perceptual disturbances (*36, 54, 55*).

Although ERPs have been recognised as disease biomarkers (*56*), their targeted monitoring and modulation is not in the focus of current therapeutic interventions. In real-life, ERP abnormalities in the context of alcohol use disorders would express in reaction, e.g. to hearing the sound of opening a beer bottle or seeing people drinking alcohol. In an addicted individual, such alcohol-associated stimuli exceedingly attract attention and challenge inhibitory abilities to withstand alcohol consumption (*57*). These processes also express as altered neural activity patterns that are supposed to normalise when neuromodulation is applied to improve behaviour control and decrease relapse risk. First studies using tDCS to correct neural disturbances in substance use disorders could positively affect N2 and P3 components (for review see (*58*)). In our study, implant-driven direct cortical stimulation successfully enhanced the N1, P2 or N2 components at most channels, indicating improved auditory processing, sensory gating and enhanced perception of the deviant stimulus. Especially for the N1 component, this effect was more pronounced at electrodes closer to the stimulation site. Nguyen and Lin (*59*) further demonstrated that the frontal N1 might not just reflect passive sensory processing but also indicates motivational salience shown by increased N1 amplitudes in rats correctly responding to a reward-coupled deviant sound compared to miss responses. Modulating the N1 component could therefore influence early inhibition of stimulus-activated actions (*60*), which is relevant to addictive diseases because relapse, as previously mentioned, is triggered by drug cue-induced craving and impaired inhibitory control (*57*). Abnormalities in stimulus-locked P2 and N2 have furthermore been associated with drug users that discontinued treatment (*61, 62*), suggesting that monitoring and modulation of these components could predict and influence treatment outcome. The P3 component was not significantly influenced by stimulation here. However, the P3 is more pronounced in the context of practical tasks involving active action planning and behaviour control than under passive conditions as applied here (*46*). Upon local delivery of NTX directly onto the PFC, we observed decreased latencies of P1 and N1 components (3 µg/µl and 30 µg/µl) and increased amplitudes of N1, P2, N2 or P3 (all doses applied) at several channels pointing to improved auditory processing. Although already approved in 1984 for addiction treatment (*63*), investigations of NTX effects on electrophysiological parameters are sparse and inconclusive. EEG studies could not detect an influence of systemic NTX application on the ERPs of our interest. In an auditory oddball paradigm with social drinkers, previous oral intake of NTX significantly reduced the late negative difference at 200 – 500 ms post-stimulus indicative for an NTX-induced impaired selective attention (*54*). In contrast, a somewhat improved neural functioning by NTX was concluded from an increased language-related N4 in a semantic memory task performed by opioid addicts, long-term treated with NTX via an abdominal subcutaneous implant (*64*).

Towards autonomous identification of brain states evoked by neuromodulation and pharmacological treatment, single-trial ERP data underwent a machine learning procedure involving SWLDA. SWLDA has a strong track record in the classification of P3-related data and has again performed well here. The comparison between sham condition and a high dose of alcohol was classified correctly in 17 out of 18 sessions, with a high degree of statistical significance (*p* < .001). Interestingly, the administration of the two different doses of alcohol was accurately distinguished, supporting recent findings in which machine learning approaches revealed that ERPs correlate with individual differences in alcohol consumption behaviour in patients suffering from alcohol use disorders (*65*). Machine learning techniques using ERP components have proven to differentiate patients from healthy controls also in schizophrenia (*66*), attention deficit hyperactive disorder (*67*) and autism (*68*). Importantly, our SWLDA approach could identify animals that received direct cortical stimulation. This is crucial for the development of closed-loop neuroprosthetics, which intend to reduce side effects and increase efficiency by switching on brain stimulation only if and as long as an abnormal neural activity is detected, adaptively adjusting stimulation parameters and switching off as soon as brain activity has been normalised (*69*). Besides the NeuroPace^®^ RNS^®^ System for epilepsy, a closed-loop system utilising a machine learning approach has been successfully implemented for DBS in Parkinson’s disease to extract patient-specific cortical signals that indicate tremor-evoking movement and adjusts stimulation voltage in real-time (*70*). ERPs and machine learning can thus support diagnosis of psychiatric symptoms and contribute to developing predictions about disease progression and treatment outcomes in individual subjects, enabling a personalised and optimised therapy (*53, 71*).

Although we have successfully demonstrated the suitability of our biomedical implant to obtain and modulate prefrontal ERPs, the safety of neural implants is crucial. Immunohistochemical analysis revealed good biocompatibility of the used implant materials, while animals in the treatment group displayed some variability in their cortical morphology. Note that animals received alcohol, electrical stimulation and local delivery of NTX. Therefore we cannot tell if the effects shown are caused by a specific treatment or result from interactions of treatments. However, this might reflect a clinical scenario where patients are treated with a combination of different approaches.

Differences in immunoreactivity were predominantly observed for laminins, a family of glycoproteins that belong to the extracellular matrix and are relevant to blood-brain-barrier and neuronal functioning (*72*). Therefore, the detected decrease in expression of laminins likely affected neurons as well. Surgical interventions to implant biomedical interfaces go along with neural and vascular injury and typical foreign body reactions to the implanted devices (*73*). Here, the necessary incision of the dura mater to allow the influx of NTX solution likely induced some vascular damage causing the observed changes in laminin expression in some animals. However, both, vascular dysfunction and neuronal loss, have also been associated with oxidative stress caused by alcohol abuse (*74*). Knabbe et al. (*75*) demonstrated that even a single intoxicating exposure to alcohol has long-lasting molecular and cellular effects in the brain. Electrical stimulation is capable to increase permeability of the blood-brain barrier as well. However, this effect is transient and reversible (*76*). Studies applying DBS suggest that electrical stimulation might regulate and even reduce neuroinflammation and apoptosis, thus excerting a neuroprotective effect (*77*). Further, alcohol-induced neurodegeneration has been associated with enhanced microglial and astrocytic expression (*78*). However, we did not observe significantly enhanced GFAP and Iba1 immunoreactivity in the treatment group. Since NTX has been shown to have anti-inflammatory and neuroprotective functions counteracting drug-induced activation of astrocytes, microglia and caspase3 (*79, 80*), we conjecture that NTX might have inhibited the potential damaging effects of repeated alcohol administration.

## Limitations

With the here applied passive auditory oddball paradigm it is not possible to clarify if targeted ERP normalisation through an epicortical implant is able to re-establish lost behaviour control. This would necessitate an experimental approach requiring subjects to actively respond to or inhibit their response to certain stimuli (*81*). When applied e.g. in addiction models, stimuli can be paired with the availability and self-administration of drugs. The implant would then allow to monitor and modulate neurophysiological correlates of drug-cue reactivity and associated drug seeking and consumption behaviour. As validated animal models are available (*82*), the experimental set-up presented here can be used straightway to target a wide range of neuropsychiatric disorders where disturbed sensory processing is a characteristic feature and closely related to the clinical symptomatology, global cognitive impairments and poor functional outcomes (*83*).

Neural measurements were currently performed following stimulation and their analysis was carried out offline. For a medical device, these elements need to be combined to enable autonomous measurement, analysis and modulation of neural activity in real-time. Integration of a controllable liquid infusion unit might allow a simpler, more straightforward and precise cortical drug delivery.

The used machine learning procedures to identify different treatment conditions were chosen on the assumption that linear classifiers perform well in the classification of brain signals compared to more complex approaches such as deep neural networks as they offer the possibility of overfitting, due to relatively small data sets usually being available (*47*). However, in this scenario, the training data set is made up of trials from numerous animals. If a large cohort of animals is used, there may be enough training data to make deep learning an attractive alternative approach. In the current data set, the stimulation electrode was the only channel consistently recorded in all sessions for all animals. Increased reliability of channel recordings may be advantageous to classification, allowing more spatial information to be utilised. The current approach relies on patterns being learned in a group of animals and transferred to a different test animal. However, brain signals are known to suffer from variability (*31*). Further sessions of all conditions would allow the use of calibration trials from the test animal’s other sessions to build a better model to generalise to a new session. With data from multiple sessions of each condition, it may even be possible to build fully subject-dependent models for each animal, which may aid accuracy.

Finally, for analysis of treatment-induced tissue reactions, rats received a combination of interventions that hinder allocation of the outcomes to a specific treatment. Further biocampatibility evaluations should be performed after longer, clinically relevant implantation durations assessing persistent effects (*73*).

## Materials and Methods

### Implant design & fabrication

The neuroprosthetic devices were manufactured using the 3D bioprinter 3DDiscovery™ Evolution (regenHU Ltd., Villaz-St-Pierre, Switzerland). The design of the implants was prepared with the printers’ bioprinting software suite BioCAD™ and in G-code using a custom-developed Python (Version 3.7) based software. The implants consisted of eight recording electrodes (0.2 × 0.2 mm^2^) and one larger frontocentral electrode (1 × 1 mm^2^) used for recording and electrical stimulation of the mPFC at 3.2 mm anterior to bregma. The electrodes were arranged in a 3 × 3 matrix with a distance between adjacent electrodes of 1.5 mm in the medio-lateral direction and 2.0 mm in the rostro-caudal direction. The implants were assembled with a 100 µm thick base layer of a silicone elastomer (DOWSIL™ SE 1700, Dow Inc., Midland, USA) with holes at the position of electrode contact sites and the microchannel. A second silicone layer on top of the base layer defined the borders of interconnects, electrodes and chemotrode. A third layer of a platinum powder (chemPUR, Karlsruhe, Germany) dispersed in tri(ethylene glycol) monomethyl ether (TGME, Merck KGaA, Darmstadt, Germany) was used for the conductive interconnects and electrode contact sites. Quality of all printing and processing steps was evaluated under a stereo microscope and defects were manually corrected. After printing each of the layers, the implants were placed on a heating plate at 120°C to enable polymerisation and evaporate the dispersing solvent. Then, a drop of PDMS (SYLGARD™184, Dow Inc., Midland, USA) pre-cured for 90 s in an oven at 90°C was manually applied under a microscope at the position of each electrode and baked on a hot plate at 105°C. The application and baking was done for each electrode separately to ensure quick polymerisation. This procedure is required to prevent formation of a thin film of PDMS covering the electrode site which is otherwise formed due to accumulation of silicone under the platinum layer. The microchannel was a silicone tubing (inner Ø: 0.51 mm, 45630102, DowCorning Silastic, Freudenberg Medical Europe GmbH, Kaiserslautern, Germany) manually connected to the implant using SE1700 silicone and allowed insertion of microsyringes to inject fluidics. Each of the interconnects was manually connected to PFA-coated stainless steel microwires (Ø: 0.23 mm, 7SS-2T, Science Products GmbH, Hofheim, Germany) using silver-containing epoxy adhesive (EPO-TEK^®^ H27D, part A, Epoxy Technology Inc., Billerica, USA) diluted in EtOH. Solvent was evaporated after application of the adhesive at 90°C. Implants were insulated with a final layer of PDMS and polymerised at 90°C. The wire connection was further insulated with a thick silicone layer using DOW CORNING^®^ 734. Finally, the microwires were soldiered to a connector (TC-2506280, Conrad Electronic SE, Hirschau, Germany) and insulated with hot glue.

### Electrochemical impedance measurements

The impedances of implant electrodes *in vitro* were recorded in PBS (pH 7.4) at room temperature using a potentiostat equipped with a frequency response analyser (AUTOLAB PGSTAT204, Deutsche Metrohm Prozessanalytik GmbH & Co. KG, Filderstadt, Germany). A platinum wire served as counter electrode and an Ag/AgCl electrode as reference. Impedance measurements *in vivo* were carried out right before each ERP recording session using the Intan RHD2000 USB interface system (Intan Technologies, Los Angeles, USA). The counter electrode was a stainless steel wire (7SS-2T, Science Products GmbH, Hofheim, Germany) whose de-insulated ending was connected to a microscrew fixed into the interparietal bone.

### Animals

All investigations within this project have been approved by the ethics committees of TU Dresden and the Saxonian ministry of the interior (Landesdirektion Sachsen, TVV 58/2018). Experiments were performed in accordance with the guidelines of the Directive 2010/63/EU on the protection of animals used for scientific purposes of the European Commission with great attention to avoid suffering and to reduce number of animals used.

The study involved n = 10 adult male Wistar wildtype rats (Janvier Labs, Le Genest-Saint-Isle, France) initially housed in groups of up to four animals. After surgery, rats were housed in single cages (Makrolon®, Type III, Tecniplast Deutschland GmbH, Hohenpeißenberg, Germany) on sawdust bedding (Ssniff - Bedding 3/4 S, Altrogge, Lage, Germany) and with Bed-r’Nest material (Datesand Ltd., Bredbury, UK) as enrichment. Pelleted food (V1534-300, ssniff Spezialdiäten GmbH, Soest, Germany) and water were available ad libitum. Housing rooms were temperature (20 - 22°C) and humidity (40 - 55 %) controlled with a 12 h automatic light-dark cycle with lights on at 6.00 am. Prior to surgery, animals were habituated to the experimenter and the recording set-up through daily handling over a duration of two weeks.

### Surgery

Surgeries to implant the device were performed under subcutaneous anaesthesia with Fentanyl (0.005 mg/kg), Midazolam (2.00 mg/kg) and Medetomidinhydrochloride (0.135 mg/kg) injected into a nuchal fold. The animals’ head was fixed into a stereotactic frame via ear pins and jaw brackets. First, the skullcap and cranial suture were exposed and two microscrews were drilled into the skull: the first one in the left parietal skull bone which is later cable-connected to the implant connector and serves as reference and the second one in the right frontal skull bone serving as anchor screw to improve fixation. The skull was slowly trepaned (6.0 mm, < 1500 rpm) under constant flushing with cold PBS at -2.6 – 3.2 mm with reference to bregma. The dura mater was then carefully incised bilaterally next to the position of the microchannel outlet (2.2 mm anterior to bregma) using microscissors. The implant was placed centrally on the cortex with the stimulation electrode located at 3.2 mm anterior to bregma. Artificial dural sealant (1A:3B, 3-4680, Dow Corning, Midland, USA) was applied on the implant to close the drill hole. The external parts of the implant were fixed to the skull with dental cement (Paladur, Kulzer GmbH, Hanau, Germany) and the wound was sutured. Upon completion of surgery, anaesthesia was antagonised by subcutaneous injection of Naloxon (0.12 mg/kg BW), Flumazenil (0.2 mg/kg) and Atipamezol (0.75 mg/kg). Animals received Meloxicam (1.0 mg/kg, s.c.) as pain medication right after surgery and on the following day.

### Experimental procedures

ECoG recordings started 3 days after surgery and were performed at a sampling rate of 3 kHz using the Intan RHD2000 USB interface system with the RHD2132 amplifier chip (Intan Technologies, Los Angeles, USA), cable-connected to the implant connector. Recordings were initially performed without and thereafter – in randomised order – every 3 days following intraperitoneal injection of 1.5 or 3 g/kg EtOH (20 % v/v, 20 min prior to recording), electrical or chemical stimulation. Electrical stimulation of the mPFC through the frontocentral electrode was applied as biphasic, charge-imbalanced pulses (100 µA anodal/-80 µA cathodal, 130 Hz) for 20 min prior to recording using a computer-interfaced current generator (STG4004, MultiChannelSystems, Reutlingen, Germany). For chemical stimulation, naltrexone (Merck KGaA, Darmstadt, Germany) was dissolved in artificial cerebrospinal fluid (125 mM NaCl, 3 mM KCl, 2.5 mM CaCl_2_, 1.3 mM MgSO_4_, 1.25 mM NaH_2_PO_4_, 26 mM NaHCO_3_, 13 mM C_6_H_12_O_6_) and applied in different dosages (3, 6 or 30 µg/µl) at a volume of 1 µl via the implant’s microchannel 20 min prior to recording.

### ECoG recording set-up

Auditory stimuli to induce ERPs were generated using custom-written Matlab scripts (Version R2019a, The Mathworks Inc., Natick, MA, USA) and consisted of frequent (standards: 50 ms, 1 kHz, 70 dB SPL, 1200 times = 87 % of trials) and rare (deviants: 50 ms, 2 kHz, 80 dB SPL, 180 times = 13 % of trials) sinusoidal sounds with rise and fall times of 5 ms and with 1 s interstimulus interval. Deviant tones were interspersed with at least one standard sound avoiding that two deviants occurred successively. One animal at a time was placed in a rodent sling (Lomir Biomedical Inc., Notre-Dame-de-l’Île-Perrot, Canada) within an electrically shielded and soundproofed audiometry booth. Sound stimuli were presented in 5 blocks of 5 min via loudspeakers at a distance of 40 cm and an angle of 45° centrally above the animals’ head.

#### Data processing & analysis

Data processing was performed using the EEGLAB toolbox (*84*) (Version 2019.1) for Matlab. Initially, data were filtered offline using a 0.1 – 45 Hz bandpass FIR filter (Kaiser windowed, Kaiser β = 5.65, filter length 54330 points). Data were segmented in epochs between -100 and 700 ms relative to stimulus onset separately for standard and deviant sounds and baseline-corrected using the pre-stimulus interval (−100 ms to 0 ms) of these epochs. Artefacts were identified and excluded based on a delta criterion of 400 µV before averaging epochs for single subjects and over all animals (grand average), respectively. ERP peak latencies were identified within standard time intervals confirmed by visual inspection (P1: 30 – 75 ms, N1: 80 – 105 ms, P2: 110 – 125 ms, N2: 130 – 180 ms, P3: 200 – 500 ms). The amplitudes of the ERP components were calculated as averaged amplitudes within a time window of 10 ms around the peak latency. Amplitudes of the P2 component are N1-P2 peak-to-peak amplitudes, whereas all other components are baseline-to-peak amplitudes.

### ERP Single-Trial Classification

Filtered and artefact-free neural recording files already segmented for standard and deviant sounds, were further used for single-trial ERP classification. Procedures were performed on the data provided by the frontocentral electrode.

#### Dataset generation for treatment classification

Data were low pass filtered using a least squares linear phase anti-aliasing FIR filter with a cut-off frequency of 32 Hz and downsampled to 64 Hz. Time domain data were then extracted in the ranges of the N1 (80 – 105 ms) and P3 (200 – 500 ms) resulting in a total number of 22 feature time points per trial. Next, ERP “difference trials” for individual sessions and animals were generated through subtracting 1.) the mean response to standard stimuli from each response to a deviant stimulus and 2.) each response to a standard stimulus from the mean response to deviant stimuli Applying both of these approaches allows the generation of a more extensive training set compared to performing only one of the methods. To train the model to predict a treatment of a particular session for a given animal, all difference trials from all other animals were combined to form the training data set. Difference trials from an animal undergoing the classification procedure were excluded from the training set.

#### Feature selection and classification

Time-domain features (i.e. the voltage values recorded at each time point, after pre-processing) were used as inputs for the feature selection phase. SWLDA feature selection first involved the creation of an initial model with no features and subsequent stepwise regression performed on the training data. A regression analysis was performed on potential models during each step, including or excluding each feature in turn and producing a F-statistic with a *p*-value for each feature. Smaller p-values indicated features with the highest likelihood of being beneficial. If any feature not already in the model had a *p*-value below the entry threshold of 0.05, then the feature with the smallest *p*-value would be added to the model. If no features were added, but any feature currently in the model now had a *p*-value above the removal threshold of 0.1, then the feature with the highest *p*-value would be removed from the model. For example, upon starting the feature selection process for a given dataset, new models would be generated containing each individual time-domain feature, and the performance of each of these models on the training data would be compared to the performance of the empty set. If we imagine that a feature achieved the lowest *p*-value at t = 250 ms, and that this *p*-value was below 0.05, the model containing only this feature would be the current model at the end of the first step. During the second step, models containing each other available feature, together with the feature at t = 250 ms, would be generated, and their performance on the training data would be compared to that of the current model. If the lowest *p*-value was achieved by a feature at t = 90 ms, and this *p*-value was below 0.05, the current model at the end of the second step would be the model containing both features at t = 90 ms and t = 250 ms. These steps continued until no features were added to, or removed from, the model. If this process failed to select any features, then the single feature with the smallest *p*-value would be selected. For the classification phase, a linear discriminant analysis model was trained and tested using the selected features. Each training trial is represented as a point in an n-dimensional space to build the model, where n is the number of selected features. A linear hyperplane is then fitted in this n-dimensional space to separate best the two sets of points representing the two classes. The class with the fewest training trials was oversampled to ensure that training occurred with an equal number of trials per class. Using this method, each difference trial in the test set from a given session was classified. In order to obtain these single-trial classifications, the test trial was represented as a point in the n-dimensional space. Depending on which side of the hyperplane it lay, it would be classified as either treatment A or treatment B. A simple majority vote was then carried out, based on the classifier outputs of all test trials in the session, to provide an overall session-level classification of which treatment had been applied. This approach was tested on one-vs-one combination for all interventions. For all treatment comparisons, each session’s actual treatments and predicted treatments were cross-tabulated to form a contingency table on which a chi-square test was performed. The classification of a given treatment comparison was considered to be statistically significant overall if the *p*-value of the chi-square statistic was less than 0.05.

### Immunohistochemistry

Analysis was performed on three groups: 1.) animals that received the surgical intervention but no implants (sham, n = 3), 2.) rats with a non-functional implant (dummy, n = 3) and 3.) animals with an active implant that received a combination of EtOH injections, electrical stimulation and NTX delivery (treatment, n = 7). After 4 weeks of implantation, rats were perfused with PBS and paraformaldehyde (PFA) and brains were extracted and stored at 4°C in PFA for 24 hours. To dehydrate the tissue, brains were kept in sucrose (30 %) at 4°C for up to one week before freezing them into methylbutan within liquid nitrogen at -40°C for 2 min. Brains were stored at -80°C until further processing. Brains were cut into slices of 40 µm thickness using a microtome and kept free-floating into anti-freeze medium at -20°C until immunohistochemical staining. Staining was performed as double-staining. From each brain, six slices were used per double-staining. Each first slice was derived from underneath the stimulation electrode at 3.2 mm anterior to bregma. Consecutive sections were always 24 sections apart covering the complete area underneath the implant. Slices were stained according to a standard staining protocol for free-floating sections. Each step preceded multiple washes in PBS. Sections were blocked in PBS containing 0.3 % Triton X-100 and 10 % serum for 2 h before incubation at 4°C overnight with primary antibodies (Table S 31) in blocking solution. The following day, slices were incubated with secondary antibodies and fluorescent dye (Table S 31) in blocking solution for 2 h before mounting on slides using Mowiol.

Fluorescent images of the brain sections were acquired with 10x magnification using the ZEISS AxioScan.Z1 Digital Slide Scanner. Image analysis was performed using the image processing suite Fiji (*85*). Thereby, images were initially background-corrected and a global threshold was applied to extract relevant objects (Fig. S 2). For each brain slice we defined a region of interest covering the entire implant width and all cortical layers up to a depth of 2 mm. The density of immunostaining (percentage of stained area per ROI, counts/mm^2^) and mean fluorescence of the six brain slices per staining were averaged to represent an animals cortical immunoreactivity to the implant and treatments.

## Statistical analysis

Statistical analysis was carried out using SPSS (Version 25, IBM Corp., Armonk, NY, USA). Initial ERP data of untreated animals underwent a one-sample *t*-test of the difference curve (deviant minus standard) against zero value. Treatment-induced modulations of neural activity were analysed applying two-tailed paired *t*-tests (*α* = 0.05) for each treatment vs. sham condition. The resulting *p*-values were corrected post-hoc for multiple comparisons (accounting for false positives amongst the 9 channels) using the Benjamini-Hochberg procedure with a threshold of 5 % and additionally reported as FDR-adjusted *p*-values. Effect sizes were calculated using Cohen’s *d* with differences of means divided by their standard deviation. Statistical analysis of immunohistochemical investigations involved a one-way ANOVA and Holm-Sidak post-hoc testing comparing numbers of stained objects, percentage of total area stained and fluorescence intensities between sham-operated animals, rats receiving an implant dummy and treatment conditions. Effect sizes were calculated using Cohen’s *f*.

## Acknowledgments

We acknowledge use of the Microstructure Facility of the BIOTEC at TU Dresden (partly funded by the State of Saxony and the European Fund for Regional Development – EFRE (100344812)) for implant fabrication and the CMCB Light Microscopy Facility for digitisation of brain tissue slides. We thank Kristin Wogan for excellent technical support.

## Funding

European Research Council (804005; IntegraBrain) (IRM)

Volkswagen Foundation (Freigeist 91 690) (IRM)

Doctoral Training Partnership scholarship from the Engineering and Physical Sciences Research Council (EPSRC) (CWirth)

Woman habilitation promotion initiative from the Medical Faculty at TU Dresden (NB)

## Author contributions

Conceptualization: BH, CWinter, IRM, NB

Methodology: BH, CWirth, DA

Software: BH, CWirth, DA

Validation: BH, CWirth, DA

Formal Analysis: BH, CWirth

Investigation: BH

Resources: IRM, NB

Data Curation: BH, CWirth

Writing – Original Draft Preparation: BH, CWirth

Writing – Review & Editing: BH, CWirth, DA, LM, CWinter, MA, IRM, NB

Visualization: BH, CWirth, DA, IRM, NB

Supervision: IRM, NB

Project administration: BH, DA, IRM, NB

Funding Acquisition: IRM, NB

## Data and materials availability

The datasets generated during and/or analysed during the current study are available from the corresponding authors on reasonable request.

## Supplementary Materials *(not included in this preprint)*

Fig. S 1 to S 5

Table S 1 to S 31

## References

1. W.-J. Gao, H.-X. Wang M. A., Y.-C. Li, in When Things Go Wrong - Diseases and Disorders of the Human Brain, T. Mantamadiotis, Ed. (InTech, 2012; http://www.intechopen.com/books/when-things-go-wrong-diseases-and-disorders-of-the-human-brain/development-of-prefrontal-cortex-and-mental-illness).

2. P. S. Olofsson, C. Bouton, Bioelectronic medicine: an unexpected path to new therapies. J Intern Med. 286, 237–239 (2019).

3. L. Agenagnew, C. kassaw, The Lifetime Prevalence and Factors Associated with Relapse Among Mentally Ill Patients at Jimma University Medical Center, Ethiopia: Cross Sectional Study. J. Psychosoc. Rehabil. Ment. Health. 7, 211–220 (2020).

4. P. Limousin, T. Foltynie, Long-term outcomes of deep brain stimulation in Parkinson disease. Nat Rev Neurol. 15, 234–242 (2019).

5. A. Merkl, G.-H. Schneider, T. Schönecker, S. Aust, K.-P. Kühl, A. Kupsch, A. A. Kühn, M. Bajbouj, Antidepressant effects after short-term and chronic stimulation of the subgenual cingulate gyrus in treatment-resistant depression. Experimental Neurology. 249, 160–168 (2013).

6. J. Kuhn, T. O. J. Gründler, R. Bauer, W. Huff, A. G. Fischer, D. Lenartz, M. Maarouf, C. Bührle, J. Klosterkötter, M. Ullsperger, V. Sturm, Successful deep brain stimulation of the nucleus accumbens in severe alcohol dependence is associated with changed performance monitoring. Addict Biol. 16, 620–623 (2011).

7. S. Ge, Y. Chen, N. Li, L. Qu, Y. Li, J. Jing, X. Wang, J. Wang, X. Wang, Deep Brain Stimulation of Nucleus Accumbens for Methamphetamine Addiction: Two Case Reports. World Neurosurgery. 122, 512–517 (2019).

8. V. Sturm, D. Lenartz, A. Koulousakis, H. Treuer, K. Herholz, J. C. Klein, J. Klosterkötter, The nucleus accumbens: a target for deep brain stimulation in obsessive–compulsive-and anxiety-disorders. Journal of Chemical Neuroanatomy. 26, 293–299 (2003).

9. C. Buhmann, T. Huckhagel, K. Engel, A. Gulberti, U. Hidding, M. Poetter-Nerger, I. Goerendt, P. Ludewig, H. Braass, C. Choe, K. Krajewski, C. Oehlwein, K. Mittmann, A. K. Engel, C. Gerloff, M. Westphal, J. A. Köppen, C. K. E. Moll, W. Hamel, Adverse events in deep brain stimulation: A retrospective long-term analysis of neurological, psychiatric and other occurrences. PLoS ONE. 12, e0178984 (2017).

10. A. Bastani, S. Jaberzadeh, Does anodal transcranial direct current stimulation enhance excitability of the motor cortex and motor function in healthy individuals and subjects with stroke: A systematic review and meta-analysis. Clinical Neurophysiology. 123, 644–657 (2012).

11. P. S. Boggio, S. P. Rigonatti, R. B. Ribeiro, M. L. Myczkowski, M. A. Nitsche, A. Pascual-Leone, F. Fregni, A randomized, double-blind clinical trial on the efficacy of cortical direct current stimulation for the treatment of major depression. International Journal of Neuropsychopharmacology. 11, 249–254 (2008).

12. J. S. Gomes, A. P. Trevizol, D. V. Ducos, A. Gadelha, B. B. Ortiz, A. O. Fonseca, H. T. Akiba, C. C. Azevedo, L. S. P. Guimaraes, P. Shiozawa, Q. Cordeiro, A. Lacerda, A. M. Dias, Effects of transcranial direct current stimulation on working memory and negative symptoms in schizophrenia: a phase II randomized sham-controlled trial. Schizophrenia Research: Cognition. 12, 20–28 (2018).

13. S. Song, A. Zilverstand, W. Gui, H. Li, X. Zhou, Effects of single-session versus multi-session non-invasive brain stimulation on craving and consumption in individuals with drug addiction, eating disorders or obesity: A meta-analysis. Brain Stimulation. 12, 606–618 (2019).

14. A.-L. Herrera-Melendez, M. Bajbouj, S. Aust, Application of Transcranial Direct Current Stimulation in Psychiatry. Neuropsychobiology. 79, 372–383 (2020).

15. A. Karabanov, A. Thielscher, H. R. Siebner, Transcranial brain stimulation: closing the loop between brain and stimulation. Current Opinion in Neurology. 29, 397–404 (2016).

16. M. Vöröslakos, Y. Takeuchi, K. Brinyiczki, T. Zombori, A. Oliva, A. Fernández-Ruiz, G. Kozák, Z. T. Kincses, B. Iványi, G. Buzsáki, A. Berényi, Direct effects of transcranial electric stimulation on brain circuits in rats and humans. Nat Commun. 9, 483 (2018).

17. D. Afanasenkau, D. Kalinina, V. Lyakhovetskii, C. Tondera, O. Gorsky, S. Moosavi, N. Pavlova, N. Merkulyeva, A. V. Kalueff, I. R. Minev, P. Musienko, Rapid prototyping of soft bioelectronic implants for use as neuromuscular interfaces. Nat Biomed Eng (2020), doi:https://doi.org/10.1038/s41551-020-00615-7.

18. I. R. Minev, P. Musienko, A. Hirsch, Q. Barraud, N. Wenger, E. M. Moraud, J. Gandar, M. Capogrosso, T. Milekovic, L. Asboth, R. F. Torres, N. Vachicouras, Q. Liu, N. Pavlova, S. Duis, A. Larmagnac, J. Voros, S. Micera, Z. Suo, G. Courtine, S. P. Lacour, Electronic dura mater for long-term multimodal neural interfaces. Science. 347, 159–163 (2015).

19. M. Athanasiadis, A. Pak, D. Afanasenkau, I. R. Minev, Direct Writing of Elastic Fibers with Optical, Electrical, and Microfluidic Functionality. Adv. Mater. Technol. 4, 1800659 (2019).

20. A. D. Degenhart, J. Eles, R. Dum, J. L. Mischel, I. Smalianchuk, B. Endler, R. C. Ashmore, E. C. Tyler-Kabara, N. G. Hatsopoulos, W. Wang, A. P. Batista, X. T. Cui, Histological evaluation of a chronically-implanted electrocorticographic electrode grid in a non-human primate. J. Neural Eng. 13, 046019 (2016).

21. E. C. Leuthardt, K. J. Miller, G. Schalk, R. P. N. Rao, J. G. Ojemann, Electrocorticography-Based Brain Computer Interface—The Seattle Experience. IEEE Trans. Neural Syst. Rehabil. Eng. 14, 194–198 (2006).

22. D. J. Caldwell, J. G. Ojemann, R. P. N. Rao, Direct Electrical Stimulation in Electrocorticographic Brain–Computer Interfaces: Enabling Technologies for Input to Cortex. Front. Neurosci. 13, 804 (2019).

23. B. Z. Mahon, M. Miozzo, W. H. Pilcher, Direct electrical stimulation mapping of cognitive functions in the human brain. Cognitive Neuropsychology. 36, 97–102 (2019).

24. F. T. Sun, M. J. Morrell, The RNS System: responsive cortical stimulation for the treatment of refractory partial epilepsy. Expert Review of Medical Devices. 11, 563–572 (2014).

25. J.-M. Fellous, G. Sapiro, A. Rossi, H. Mayberg, M. Ferrante, Explainable Artificial Intelligence for Neuroscience: Behavioral Neurostimulation. Front. Neurosci. 13, 1346 (2019).

26. E. M. Sokhadze, M. F. Casanova, E. L. Casanova, E. Lamina, D. P. Kelly, I. Khachidze, Event-related Potentials (ERP) in Cognitive Neuroscience Research and Applications. NR. 4, 14–27 (2017).

27. E. S. Kappenman, S. J. Luck, ERP Components: The Ups and Downs of Brainwave Recordings (Oxford University Press, 2011; http://oxfordhandbooks.com/view/10.1093/oxfordhb/9780195374148.001.0001/oxfordhb-9780195374148-e-001).

28. M. Lijffijt, S. D. Lane, S. L. Meier, N. N. Boutros, S. Burroughs, J. L. Steinberg, F. Gerard Moeller, A. C. Swann, P50, N100, and P200 sensory gating: Relationships with behavioral inhibition, attention, and working memory. Psychophysiology. 46, 1059–1068 (2009).

29. S. Campanella, O. Pogarell, N. Boutros, Event-Related Potentials in Substance Use Disorders: A Narrative Review Based on Articles from 1984 to 2012. Clin EEG Neurosci. 45, 67–76 (2014).

30. J. R. Folstein, C. Van Petten, Influence of cognitive control and mismatch on the N2 component of the ERP: A review. Psychophysiology. 45 (2008), doi:10.1111/j.1469-8986.2007.00602.x.

31. J. Polich, Updating P300: An integrative theory of P3a and P3b. Clinical Neurophysiology. 118, 2128–2148 (2007).

32. R. T. Knight, W. Richard Staines, D. Swick, L. L. Chao, Prefrontal cortex regulates inhibition and excitation in distributed neural networks. Acta Psychologica. 101, 159–178 (1999).

33. C. L. Ehlers, ERP responses to ethanol and diazepam administration in squirrel monkeys. Alcohol. 5, 315–320 (1988).

34. H. L. Cohen, J. Ji, D. B. Chorlian, H. Begleiter, B. Porjesz, Alcohol-related ERP changes recorded from different modalities: a topographic analysis. Alcohol Clin Exp Res. 26, 303–317 (2002).

35. J. Marco, L. Fuentemilla, C. Grau, Auditory sensory gating deficit in abstinent chronic alcoholics. Neuroscience Letters. 375, 174–177 (2005).

36. C. L. Ehlers, A. Desikan, D. N. Wills, Event-Related Potential Responses to the Acute and Chronic Effects of Alcohol in Adolescent and Adult Wistar Rats. Alcohol Clin Exp Res. 38, 749–759 (2014).

37. G. Petit, A. Cimochowska, C. Cevallos, G. Cheron, C. Kornreich, C. Hanak, E. Schroder, P. Verbanck, S. Campanella, Reduced processing of alcohol cues predicts abstinence in recently detoxified alcoholic patients in a three-month follow up period: An ERP study. Behavioural Brain Research. 282, 84–94 (2015).

38. M. Luijten, M. Kleinjan, I. H. A. Franken, Event-related potentials reflecting smoking cue reactivity and cognitive control as predictors of smoking relapse and resumption. Psychopharmacology. 233, 2857–2868 (2016).

39. J. L. Voss, K. A. Paller, in Learning and Memory: A Comprehensive Reference (Elsevier, 2017; https://linkinghub.elsevier.com/retrieve/pii/B9780128093245210705), pp. 81–98.

40. D. J. Krusienski, J. J. Shih, in 2010 Annual International Conference of the IEEE Engineering in Medicine and Biology (IEEE, Buenos Aires, 2010; http://ieeexplore.ieee.org/document/5627603/), pp. 6019–6022.

41. B. E. Mouthaan, M. A. van ‘t Klooster, D. Keizer, G. J. Hebbink, F. S. S. Leijten, C. H. Ferrier, M. J. A. M. van Putten, M. Zijlmans, G. J. M. Huiskamp, Single Pulse Electrical Stimulation to identify epileptogenic cortex: Clinical information obtained from early evoked responses. Clinical Neurophysiology. 127, 1088–1098 (2016).

42. A. L. Benabid, T. Costecalde, A. Eliseyev, G. Charvet, A. Verney, S. Karakas, M. Foerster, A. Lambert, B. Morinière, N. Abroug, M.-C. Schaeffer, A. Moly, F. Sauter-Starace, D. Ratel, C. Moro, N. Torres-Martinez, L. Langar, M. Oddoux, M. Polosan, S. Pezzani, V. Auboiroux, T. Aksenova, C. Mestais, S. Chabardes, An exoskeleton controlled by an epidural wireless brain–machine interface in a tetraplegic patient: a proof-of-concept demonstration. The Lancet Neurology. 18, 1112–1122 (2019).

43. M. J. Vansteensel, E. G. M. Pels, M. G. Bleichner, M. P. Branco, T. Denison, Z. V. Freudenburg, P. Gosselaar, S. Leinders, T. H. Ottens, M. A. Van Den Boom, P. C. Van Rijen, E. J. Aarnoutse, N. F. Ramsey, Fully Implanted Brain–Computer Interface in a Locked-In Patient with ALS. N Engl J Med. 375, 2060–2066 (2016).

44. D. R. Merrill, M. Bikson, J. G. R. Jefferys, Electrical stimulation of excitable tissue: design of efficacious and safe protocols. Journal of Neuroscience Methods. 141, 171–198 (2005).

45. C. S. Hendershot, J. D. Wardell, A. V. Samokhvalov, J. Rehm, Effects of naltrexone on alcohol self-administration and craving: meta-analysis of human laboratory studies: Naltrexone human lab studies. Addiction Biology. 22, 1515–1527 (2017).

46. E. Wronka, J. Kaiser, A. M. L. Coenen, The auditory P3 from passive and active three-stimulus oddball paradigm. Acta Neurobiologiae Experimentalis. 68, 362–372 (2008).

47. F. Lotte, M. Congedo, A. Lécuyer, F. Lamarche, B. Arnaldi, A review of classification algorithms for EEG-based brain–computer interfaces. J. Neural Eng. 4, R1–R13 (2007).

48. D. J. Krusienski, E. W. Sellers, F. Cabestaing, S. Bayoudh, D. J. McFarland, T. M. Vaughan, J. R. Wolpaw, A comparison of classification techniques for the P300 Speller. J. Neural Eng. 3, 299–305 (2006).

49. C. Wirth, J. Toth, M. Arvaneh, “You Have Reached Your Destination”: A Single Trial EEG Classification Study. Front. Neurosci. 14, 66 (2020).

50. A. A. Schendel, M. W. Nonte, C. Vokoun, T. J. Richner, S. K. Brodnick, F. Atry, S. Frye, P. Bostrom, R. Pashaie, S. Thongpang, K. W. Eliceiri, J. C. Williams, The effect of micro-ECoG substrate footprint on the meningeal tissue response. J. Neural Eng. 11, 046011 (2014).

51. A. Stiller, J. Usoro, J. Lawson, B. Araya, M. González-González, V. Danda, W. Voit, B. Black, J. Pancrazio, Mechanically Robust, Softening Shape Memory Polymer Probes for Intracortical Recording. Micromachines. 11, 619 (2020).

52. M. D. Johnson, K. J. Otto, D. R. Kipke, Repeated Voltage Biasing Improves Unit Recordings by Reducing Resistive Tissue Impedances. IEEE Trans. Neural Syst. Rehabil. Eng. 13, 160–165 (2005).

53. S. Campanella, Why it is time to develop the use of cognitive event-related potentials in the treatment of psychiatric diseases. NDT, 1835 (2013).

54. I. P. Jääskeläinen, J. Hirvonen, T. Kujala, K. Alho, C. J. P. Eriksson, A. Lehtokoski, E. Pekkonen, J. D. Sinclair, H. Yabe, R. Näätänen, P. Sillanaukee, Effects of naltrexone and ethanol on auditory event-related brain potentials. Alcohol. 15, 105–111 (1998).

55. D. C. Gooding, K. Gjini, S. A. Burroughs, N. N. Boutros, The association between psychosis proneness and sensory gating in cocaine-dependent patients and healthy controls. Psychiatry Research. 210, 1092–1100 (2013).

56. S. Campanella, C. Colin, Event-related potentials and biomarkers of psychiatric diseases: the necessity to adopt and develop multi-site guidelines. Front. Behav. Neurosci. 8 (2014), doi:10.3389/fnbeh.2014.00428.

57. S. Campanella, Neurocognitive rehabilitation for addiction medicine, in Progress in Brain Research (Elsevier, 2016; https://linkinghub.elsevier.com/retrieve/pii/S007961231500120X), vol. 224, pp. 85–103.

58. B. Habelt, M. Arvaneh, N. Bernhardt, I. Minev, Biomarkers and neuromodulation techniques in substance use disorders. Bioelectron Med. 6, 4 (2020).

59. D. P. Nguyen, S.-C. Lin, A frontal cortex event-related potential driven by the basal forebrain. Elife. 3, e02148 (2014).

60. J. B. Debruille, M. Touzel, J. Segal, C. Snidal, L. Renoult, A Central Component of the N1 Event-Related Brain Potential Could Index the Early and Automatic Inhibition of the Actions Systematically Activated by Objects. Front. Behav. Neurosci. 13, 95 (2019).

61. B. C. Fink, V. R. Steele, M. J. Maurer, S. J. Fede, V. D. Calhoun, K. A. Kiehl, Brain potentials predict substance abuse treatment completion in a prison sample. Brain Behav. 6 (2016), doi:10.1002/brb3.501.

62. V. R. Steele, B. C. Fink, J. M. Maurer, M. R. Arbabshirani, C. H. Wilber, A. J. Jaffe, A. Sidz, G. D. Pearlson, V. D. Calhoun, V. P. Clark, K. A. Kiehl, Brain Potentials Measured During a Go/NoGo Task Predict Completion of Substance Abuse Treatment. Biological Psychiatry. 76, 75–83 (2014).

63. U.S. Department of Health and Human Services, in Incorporating Alcohol Pharmacotherapies Into Medical Practice (Substance Abuse and Mental Health Services Administration, Rockville, MD, USA, 2009; https://www.ncbi.nlm.nih.gov/books/NBK64042/), vol. 49.

64. H. Sheng-xi, Y. Long-chuan, C. Qing, W. Dong-mei, H. Shu, J. Shao-wei, Effect of long-term sustained release naltrexone on semantic recognition of opioid addicts. Journal of Clinical Rehabilitative Tissue Engineering Research. 13, 1573–1576 (2009).

65. L. O’Halloran, L. M. Rueda-Delgado, L. Jollans, Z. Cao, R. Boyle, C. Vaughan, P. Coey, R. Whelan, Inhibitory-control event-related potentials correlate with individual differences in alcohol use. Addiction Biology. 25 (2020), doi:10.1111/adb.12729.

66. A. H. Neuhaus, F. C. Popescu, J. A. Bates, T. E. Goldberg, A. K. Malhotra, Single-subject classification of schizophrenia using event-related potentials obtained during auditory and visual oddball paradigms. Eur Arch Psychiatry Clin Neurosci. 263, 241–247 (2013).

67. A. Mueller, G. Candrian, J. D. Kropotov, V. A. Ponomarev, G.-M. Baschera, Classification of ADHD patients on the basis of independent ERP components using a machine learning system. Nonlinear Biomed Phys. 4 Suppl 1, S1 (2010).

68. J. Eldridge, A. E. Lane, M. Belkin, S. Dennis, Robust features for the automatic identification of autism spectrum disorder in children. J Neurodev Disord. 6, 12 (2014).

69. M. Parastarfeizabadi, A. Z. Kouzani, Advances in closed-loop deep brain stimulation devices. J NeuroEngineering Rehabil. 14, 79 (2017).

70. B. Houston, M. Thompson, A. Ko, H. Chizeck, A machine-learning approach to volitional control of a closed-loop deep brain stimulation system. J. Neural Eng. 16, 016004 (2019).

71. A. Thieme, D. Belgrave, G. Doherty, Machine Learning in Mental Health: A Systematic Review of the HCI Literature to Support the Development of Effective and Implementable ML Systems. ACM Trans. Comput.-Hum. Interact. 27, 1–53 (2020).

72. A. W. Lasek, Effects of Ethanol on Brain Extracellular Matrix: Implications for Alcohol Use Disorder. Alcohol. Clin. Exp. Res. 40, 2030–2042 (2016).

73. D. Prodanov, J. Delbeke, Mechanical and Biological Interactions of Implants with the Brain and Their Impact on Implant Design. Front. Neurosci. 10 (2016), doi:10.3389/fnins.2016.00011.

74. S. Alikunju, P. M. Abdul Muneer, Y. Zhang, A. M. Szlachetka, J. Haorah, The inflammatory footprints of alcohol-induced oxidative damage in neurovascular components. Brain, Behavior, and Immunity. 25, S129–S136 (2011).

75. J. Knabbe, J. Protzmann, N. Schneider, D. Dannehl, M. Berger, S. Wei, C. Strahle, A. Jaiswal, S. Lugani, H. Zheng, M. Krüger, K. Rohr, R. Spanagel, H. Scholz, A. Bilbao, M. Engelhardt, S. B. Cambridge, “Single-dose ethanol intoxication causes acute and lasting neuronal changes in the brain” (preprint, Neuroscience, 2020),, doi:10.1101/2020.09.09.289256.

76. D. W. Shin, J. Fan, E. Luu, W. Khalid, Y. Xia, N. Khadka, M. Bikson, B. M. Fu, In Vivo Modulation of the Blood–Brain Barrier Permeability by Transcranial Direct Current Stimulation (tDCS). Ann Biomed Eng. 48, 1256–1270 (2020).

77. C. McKinnon, P. Gros, D. J. Lee, C. Hamani, A. M. Lozano, L. V. Kalia, S. K. Kalia, Deep brain stimulation: potential for neuroprotection. Ann Clin Transl Neurol. 6, 174–185 (2019).

78. M. Saito, G. Chakraborty, M. Hui, K. Masiello, M. Saito, Ethanol-Induced Neurodegeneration and Glial Activation in the Developing Brain. Brain Sciences. 6 (2016), doi:10.3390/brainsci6030031.

79. M. R. Hutchinson, Y. Shavit, P. M. Grace, K. C. Rice, S. F. Maier, L. R. Watkins, Exploring the neuroimmunopharmacology of opioids: an integrative review of mechanisms of central immune signaling and their implications for opioid analgesia. Pharmacol. Rev. 63, 772–810 (2011).

80. L. Qin, F. T. Crews, Chronic ethanol increases systemic TLR3 agonist-induced neuroinflammation and neurodegeneration. J Neuroinflammation. 9, 130 (2012).

81. M. Kutas, M. Kiang, K. Sweeney, in The Handbook of the Neuropsychology of Language, M. Faust, Ed. (Wiley-Blackwell, Oxford, UK, 2012; http://doi.wiley.com/10.1002/9781118432501.ch26), pp. 543–564.

82. E. J. Nestler, S. E. Hyman, Animal models of neuropsychiatric disorders. Nat Neurosci. 13, 1161–1169 (2010).

83. I. S. Ramsay, M.-P. Schallmo, B. Biagianti, M. Fisher, S. Vinogradov, S. R. Sponheim, Deficits in Auditory and Visual Sensory Discrimination Reflect a Genetic Liability for Psychosis and Predict Disruptions in Global Cognitive Functioning. Front. Psychiatry. 11, 638 (2020).

84. A. Delorme, S. Makeig, EEGLAB: an open source toolbox for analysis of single-trial EEG dynamics including independent component analysis. Journal of Neuroscience Methods. 134, 9–21 (2004).

85. J. Schindelin, I. Arganda-Carreras, E. Frise, V. Kaynig, M. Longair, T. Pietzsch, S. Preibisch, C. Rueden, S. Saalfeld, B. Schmid, J.-Y. Tinevez, D. J. White, V. Hartenstein, K. Eliceiri, P. Tomancak, A. Cardona, Fiji: an open-source platform for biological-image analysis. Nat Methods. 9, 676–682 (2012).

